# Neural representation of spatial and non-spatial auditory attention in EEG signals

**DOI:** 10.1101/2023.07.13.548897

**Authors:** Winko W. An, Abigail Noyce, Alexander Pei, Barbara Shinn-Cunningham

**Author notes:** Corresponding author. Baker Hall 254G, Carnegie Mellon University, 5000 Forbes Avenue, Pittsburgh, PA, 15213, USA. Email addresses:* *(Winko W. An)*, *(Abigail Noyce)*, *(Alexander Pei)*, *(Barbara Shinn-Cunningham).

## Abstract

Neural representation, capturing the content and format of encoded information, provides insight into the internal states of neural units. Studies of neural representation contrast with studies of neural processes, which focus on how one neural unit influences another. Representational similarity analysis (RSA), a multivariate analysis approach, has been used in previous studies to explore the neural representation of object categories in various neuroimaging modalities. In this study, we employed RSA to examine the neural representation of executive function. We designed an experiment involving a rich set of conditions where participants engaged in an auditory task requiring either spatial or non-spatial attention. We extracted representational features from their electroencephalography (EEG) scalp voltage and alpha power and compared these features with ideal conceptual models representing perfect categorization of different attentional states. The results demonstrate the feasibility of investigating internal cognitive states using RSA. Specifically, we identified time intervals during which attentional state contrasts, such as differences between attention types or locations, manifested in the measured neural responses from scalp voltage and alpha power distributions.

## 1. Introduction

Electroencephalography (EEG) requires only simple setup, costs very little, and provides high temporal resolution, making it a popular tool to study the dynamics of cognitive functions in the human brain. EEG monitors neural activity by sampling the electrical potential at the scalp at a very high rate. It carries rich spectrotemporal information and follows the fast dynamics of brain response to sensory inputs. By measuring evoked event-related potentials (ERPs) and induced neural oscillations, neuroscientists have used EEG to characterize how cognitive functions, such as top-down attentional control, modulate the amplitude and/or phase of these neural signatures (Herrmann and Knight, 2001; Siegel et al., 2012). Such traditional studies typically analyze different electrical measures (e.g., amplitude, latency, power, etc.) at a single electrode or over a small region-of-interest (ROI) in sensor space. This type of univariate analysis allows researchers to make inferences about the *neural processes* underpinning cognitive functions. Recently, with advances in machine learning algorithms, multivariate analysis has become another popular approach to studying neural processes reflected in high-dimensional measurements. This type of analysis can be applied to EEG signals by grouping together signals from multiple electrodes and/or time points to form multivariate features. These high-dimensional features are then used to train and test classifiers (usually for binary classification); when appropriately cross-validated, the resulting classification accuracy serves as a powerful proxy for quantifying differences in brain responses across experimental conditions. Typically, however, multivariate analyses have compared neural responses across only a small number of conditions.

Over the past decade, more and more studies have begun to use multi-variate analysis of high-dimensional brain data to explore the *neural representation* of different forms of information. The term *neural representation* refers specifically to the content and format of information that a neural/cognitive unit encodes stably: a neural representation reflects internal states of the underlying neural unit (Freund et al., 2021). This definition contrasts with the term *neural process*, which typically refers to how one neural/cognitive function acts on another system or outcome (e.g., how “attention” modulates the “ERP amplitude”).

*Representational similarity analysis* (RSA) provides a way to study neural representation for experiments employing a condition-rich design (Kriegeskorte et al., 2008). In this framework, one first computes a set of *representational dissimilarity matrices* (RDMs) from multivariate neuroimaging data (e.g., EEG): each matrix has one row and one column for each condition, with each entry in the matrix quantifying the difference between the neural measures in response to the corresponding pair of conditions (e.g., derived using appropriate classification or neural distance metrics). The similarity structure of the resulting RDMs provides an abstract representation of the information captured by the neural metrics used to build the neural RDM and encoded by the brain. One can then make inferences about the underlying neural representation by comparing these neuroimaging RDMs to idealized model RDMs, whose structure can be built from specific hypotheses about how the neural representation should differ across conditions given the experimental design. Specifically, when the neural RDM matches the structure of an idealized model RDM, it provides support for the specific hypothesis generating the model RDM.

The RSA approach has been widely used to study dynamics in sensory processing. For example, Cichy et al. (2014) designed an experiment in which they presented participants with a wide range of visual objects of different categories and collected both magnetoencephalography (MEG) and functional magnetic resonance imaging (fMRI). From the MEG signals, they calculated RDMs at each time point to reveal how the encoded visual information evolved as a function of time. From the fMRI responses, they calculated RDMs at each voxel to reveal where in the brain different representations occurred. Critically, they estimated the information flow in the brain during visual object recognition by comparing fMRI and MEG RDMs to find where in the brain (from fMRI RDMs) information matched that occurring at different time points (from MEG RDMs). RSA has also been applied to EEG signals in other domains to trace dynamics of various neural processes, including audiovisual integration (Cecere et al., 2017), visual object processing (Kaneshiro et al., 2015) and memory (Sommer et al., 2019).

These studies share a common trait: by using a range of different sensory stimuli across conditions, they explored how different representations of stimuli emerge over time and / or where these representations arise. However, none explored the internal cognitive state of the observer (e.g., examining how task goals change the information flow for the same types of sensory inputs). Theoretically, though, RSA can also be applied to study cognitive control. The abstract nature of similarity structures allow RSA the flexibility to fit various cognitive tasks, while conceptual model RDMs can explicitly test whether the resulting neural representations match those predicted by specific hypotheses. This makes inferences straightforward (Freund et al., 2021).

In this study, we deploy RSA to investigate the neural representation of attentional states during auditory processing. We designed an experiment in which input auditory signals were statistically identical, then varied whether listeners engaged spatial or non-spatial auditory attention. We used features in EEG scalp voltage topography and its alpha power topography for neural decoding to understand the degree to which the listener’s internal attentional states differed across these two attentional conditions. Based on previous work, we expected the different forms of attention to differentially engage a fronto-parietal network specialized for processing visuo-spatial information (Michalka et al., 2015, 2016); further, we expected neural oscillations over parietal cortex to differ, depending on how auditory attention was focused (e.g., see Deng et al. (2020)). This approach allowed us to build specific model RDMs against which we could compare the observed neural RDMs. We then compared the neural RDMs to our model RDMs to reveal the kind of information being encoded at each time point during the experiment.

## 2. Materials and methods

All study procedures were approved by the Institution Review Board of Boston University.

### 2.1. Participants

Thirty adults (19 – 44 years old, 14 women) participated in this study. No participant reported hearing loss or any history of neurological disorders. All participants gave written informed consent, and were paid for their time.

### 2.2. Stimuli and task

Participants listened for the identity of a target syllable (/ba/, /da/, or /ga/) presented amidst three distractor syllables. The syllables were spatialized using a generic head-related transfer function (Gardner and Martin, 1994) that imbued each syllable with spatial cues (interaural time differences, interaural level differences, and spectral cues) associated with one of five locations in azimuth (in the horizontal plane of the ears): 90° from the left (L90), 30° from the left (L30), center, 30° from the right (R30), or 90° from the right (R90) (Figure 1a).

**Figure 1:**
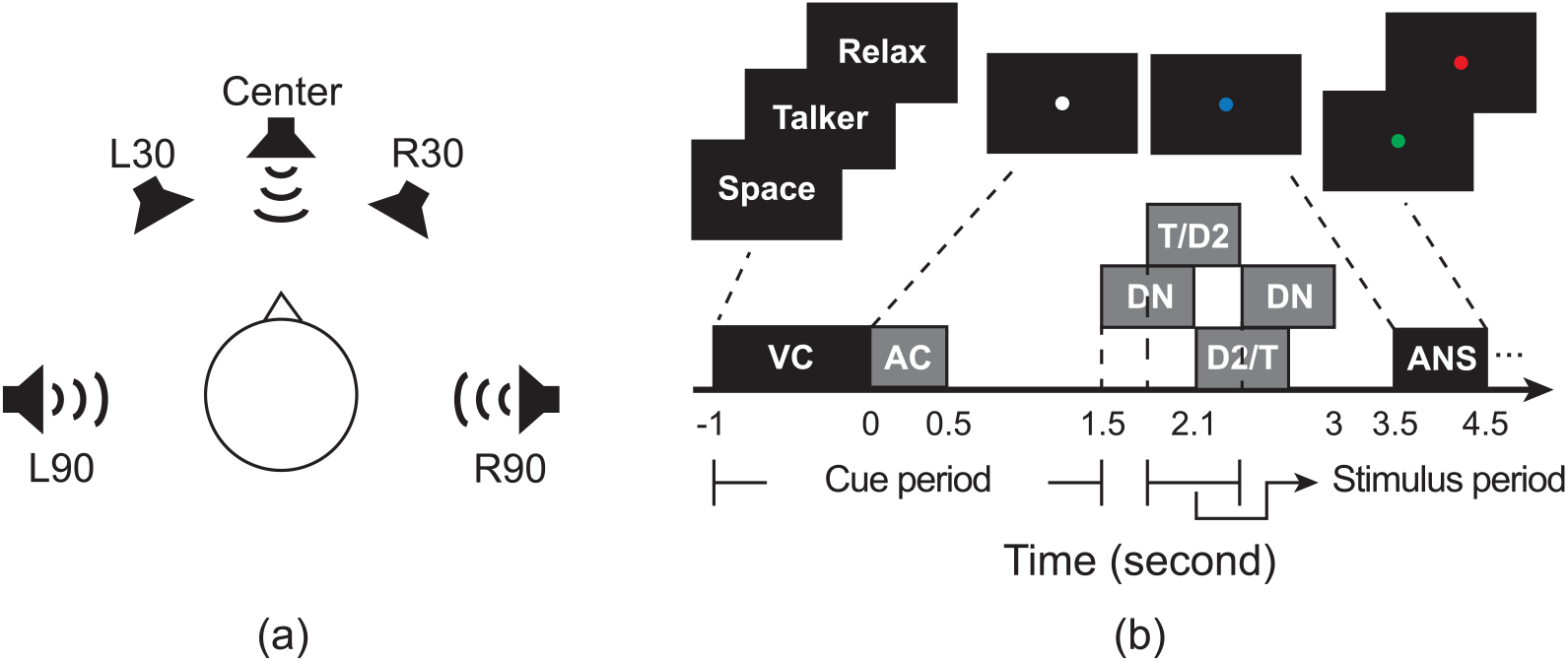
Experimental task. (a) Spoken syllables were spatialized to center, 30° left (L30) or right (R30), and 90° left (L90) or right (R90), always in the horizontal plane. Depending on the trial type, listeners were instructed to attend to the target based on either its location or the talker producing it, or to ”relax” and not attend. (b) Illustration of the events within a trial for one example trial type. A visual cue (VC) indicated the type of attention required for that trial. A subsequent auditory cue (AC) then came on; this cue had the ”value” of the feature to attend (coming from a specific direction or talker that the listener had to then focus on in the subsequent syllable mixture). A 4-syllable mixture was played one second after the AC. The first and fourth syllables were always neutral distractors (DN; they come from the center with an intermediate pitch). Of the second and the third syllables, one was the target (T), and the other served as an additional distractor (D2). All syllables were 600ms long, and their onsets were 300ms apart. The participants were instructed to respond by identifying the target syllable when the fixation dot turned blue, immediately after the end of the final distractor stimulus. Feedback was given at the end of each trial; the fixation dot turned either green (correct) or red (incorrect).

Figure 1b shows the sequence of events in a trial. A visual cue (VC) instructed the listener to use either spatial attention (Space) or non-spatial attention (Talker), or to passively listen wihout performing a particular task (Relax). Next, an auditory cue (AC) (/a/) came on that had the value of the attentional feature that listeners had to use to pick out the target syllable in the upcoming mixture of syllables for that trial. Specifically, in Space trials, the AC came from the target location (either L90 or R90) but was spoken in a neutral voice; in Talker trials, the AC was spoken in the target voice (one of four talkers) but came from the center (which was never the target syllable direction); in Relax trials, both features were neutral (center location, synthesized gender-neutral voice). A 4-syllable mixture was then presented. The target syllable was either the second or third syllable, spoken by one of four talkers and coming from either L90 or R90, with the appropriate target feature matching the AC. The (other) third or second syllable was a distractor that always was spoken by a different talker than the target (see Table 1). The first and fourth syllables always came from the center, from a neutral voice, and were always distractors. Unless it was a Relax trial, the listeners were instructed to report the identity of the target syllable, which required them to deploy either spatial or non-spatial attention in order to be correct. On Relax trials, listeners were asked to withhold response and make a random key press during the response period.

**Table 1:** The 21 experimental conditions used in this study. Conditions differed in their attention type, the location or gender of talker for the target (T), and the location and gender of the talker for the second distractor (D2). The “Relax” conditions used the same stimuli as in “Space” and “Talker”, but required no attention.

| Attention type | Location of T | Location of D2 | T & D2 gender | Condition # |
| --- | --- | --- | --- | --- |
| Space | L90 | L30 | different | 1 |
|  |  |  | same | 2 |
|  |  | R90 | different | 3 |
|  |  |  | same | 4 |
|  | R90 | R30 | different | 5 |
|  |  |  | same | 6 |
|  |  | L90 | different | 7 |
|  |  |  | same | 8 |

| Attention type | Gender of T | Gender of D2 | T & D2 location |  |
| --- | --- | --- | --- | --- |
| Talker | female | male | same location | 9 |
|  |  |  | different locations, same side | 10 |
|  |  |  | different sides | 11 |
|  | male | female | same location | 12 |
|  |  |  | different locations, same side | 13 |
|  |  |  | different sides | 14 |

| Attention type | Using the same stimuli as in condition # |  |  |  |
| --- | --- | --- | --- | --- |
| Relax | 1, 10, 13 |  |  | 15 |
|  | 2 |  |  | 16 |
|  | 3, 7, 11, 14 |  |  | 17 |
|  | 4, 8 |  |  | 18 |
|  | 5, 10, 13 |  |  | 19 |
|  | 6 |  |  | 20 |
|  | 9, 12 |  |  | 21 |

### 2.3. Experiment Design and Procedures

The full experiment consisted of 21 conditions (Table 1). Each condition used a different specific combination of 1) type of attention required, 2) characteristics of the target, and 3) characteristics of the distractors. Stimulus characteristics were matched between Space, Talker, and Relax conditions, so any differences in brain signals could be attributed unambiguously to differences in internal attentional state. Participants completed 36 trials of each condition for a total of 756 trials. The order of trials was shuffled and evenly divided into 12 blocks with short breaks in between for participant comfort. Testing for each subject was conducted in a single 2-hour long session over 12 blocks with brief, self-paced breaks in between blocks.

### 2.4. EEG recording and preprocessing

EEG was continuously recorded from 64 electrodes according to the standard 10/20 system. Signals were digitized at 2048 Hz using the ActiveTwo system (BioSemi B.V.), passed through a sinc windowed FIR band-pass filter (0.1 – 50 Hz) to remove slow drifts and line noise, and downsampled to 256 Hz. An independent component analysis using EEGLAB (Delorme and Makeig, 2004) allowed us to manually remove components that were eye blinks or muscle artifacts. The continuous EEG signals were then split into epochs. We isolated a preparatory attention period from −1000 – 1500 ms relative to the onset of the auditory cue (cue period), and a peristimulus attention period from −300 – 500 ms relative to the onset of the target syllable (stimulus period). In this methods-focused paper, we focus on data taken from the cue period. Epochs were baseline corrected using the average signal during the 100 ms before the onset of VC. Epochs with substantial noise after ICA (absolute amplitude over 20 *µ*V; visible muscle artifact) were manually removed from further analysis. After preprocessing, an average of 735.4 *±* 22.9 trials were retained per individual subject (min had 660, max had 756); an average of 35.0 *±* 1.4 trials were retained for each condition for each subject (min had 28, max had 36).

### 2.5. Measuring representational dissimilarity

In order to study the neural representation of attentional states, we decoded auditory attention from EEG signals. Here we show decoding for 2 features: 1) the EEG scalp voltage topography (Scalp Voltage), and 2) the alpha-band (8 – 14 Hz) power topography (Alpha Power). Figure 2 summarizes the steps in extracting neural representations for attention decoding.

**Figure 2:**
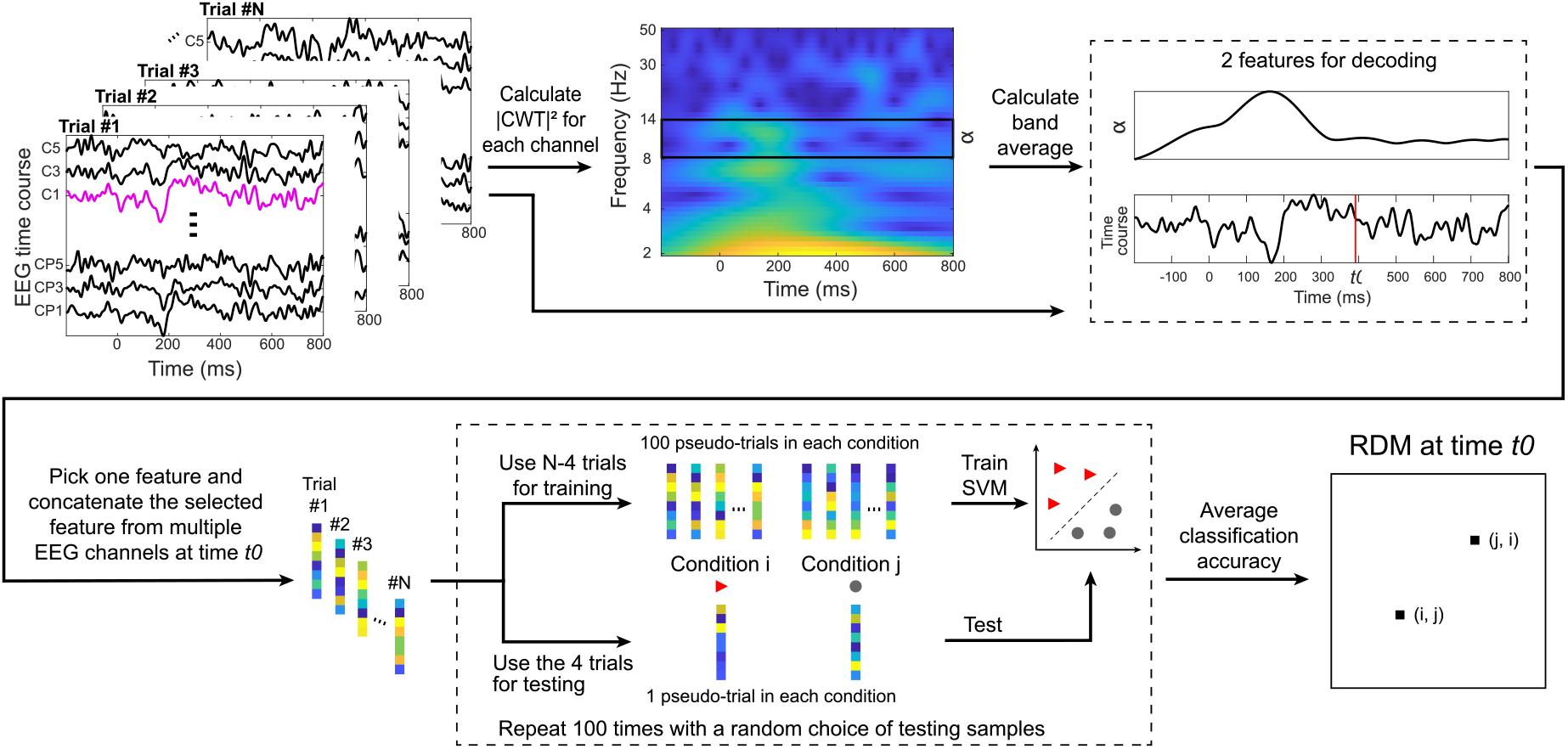
Schematic diagram of how a representational dissimilarity matrix (RDM) was derived. Either the EEG scalp voltage topography or alpha-band power topography (calculated from a continuous wavelet transform, or CWT) was used as the feature to decode attention at a particular time point. Dissimilarity between each pair of conditions was estimated through 100 iterations of training-and-test of a support vector machine (SVM) with a random choice of test samples, selected to be independent of the training data, within each iteration. The average classification accuracy was saved in an RDM at its corresponding positions. The output of this attention decoding procedure is a series of RDMs that create a time-varying function for each feature of choice (the EEG Scalp Voltage or power of a frequency band).

The two features on which we focused — the EEG Scalp Voltage and the instantaneous Alpha Power — were studied independently. For the Scalp Voltage, the feature for a subject at a time instance *t*_0_ comprised the preprocessed 64-channel EEG signal at that time point; the length of this feature vector was thus 64. For the Alpha Power, we calculated a continuous wavelet transform (CWT) for each channel and averaged across the frequencies within the alpha frequency range. The dimensionality of an Alpha Power feature vector at *t*_0_ was thus also 64.

To boost the signal-to-noise ratio in feature vectors, we adopted a bootstrapped averaging method with replacement (Grootswagers et al., 2017). In each iteration, we randomly selected 4 trials from a condition, averaged their vectors to generate a pseudotrial, and used this pseudotrial as a testing sample. Over the rest of the trials in that condition, we randomly selected four at a time and calculated their average, and repeated this process 100 times to generate 100 pseudotrials for training. This method was demonstrated to be effective in boosting the decoding accuracy (see below) when there is a real difference between conditions (Grootswagers et al., 2017).

Using the derived distributions of these pseudotrials, we calculated the representational dissimilarities at each time point between each pair of conditions. We trained a linear support vector machine (SVM) and evaluated the dissimilarity between conditions as the average decoding accuracy across 100 iterations of cross-validation. If two conditions are significantly dissimilar (e.g., a Space condition vs a Relax condition), an SVM can learn their differences from training samples and yield above-chance classification performance (i.e., *>*50%). Conversely, if two conditions utilize the same type of attention, the decoding accuracy might be close to chance.

We summarized the dissimilarities between all pairs of conditions in a representational dissimilarity matrix (RDM). Each row and column of an RDM corresponds to a condition index, and each element corresponds to the dissimilarity value between two conditions: the classification accuracy when discriminating between condition i and condition j fills both the (i, j) and (j, i) elements of the RDM. The major diagonal of an RDM, representing the difference between one condition and itself, is filled with zeros and is excluded from any analysis. For each subject, two sets of RDMs were generated from this attention decoding process, one for the Scalp Voltage and one for the Alpha Power. Because there is one RDM at each time instance, each RDM set can be represented as a function of time, summarizing the temporal dynamics of the neural representation of attention. Of course, the samples across time are not independent, given the natural frequency content of these signals and the structure imposed by the filtering and other data processing steps. To account for this temporal structure in the time-varying RDMs, we employed a permutation approach (see Section 2.7 for details) to ensure that all statistical tests accounted for temporal correlations in the underlying data.

### 2.6. Comparison to conceptual models

Because an RDM contains dissimilarity measures among all pairs of conditions, we can inspect particular submatrices to understand whether two categories of attention are represented differently at a time point (Figure 3a). For example, at moments in which focusing spatial attention produces a clear brain state that is reflected in the neural metrics, we expect entries in the RDM comparing Space-Relax conditions to carry high values reflecting large dissimilarity between the spatial-attention state and the ”no task” state, but submatrices reflecting comparisons between Space-Space conditions to carry relatively low values, reflecting the similarity of the brain responses for comparison.

**Figure 3:**
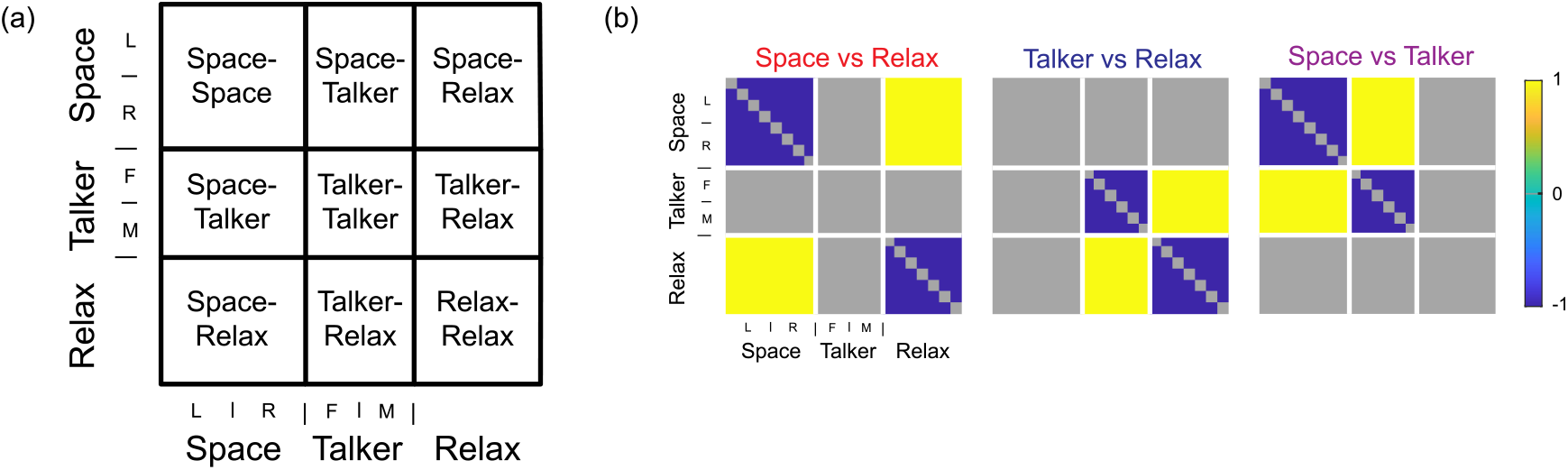
(a) A representational dissimilarity matrix (RDM) in this study can be divided into several submatrices, each of which represents a different comparison between types of internal attentional / task state. The top left portion of the RDM, Space-Space, shows the differences between conditions differing in stimuli but sharing a goal of focusing spatial attention. Similarly, the other two square submatrices along the major diagonal, Talker-Talker and Relax-Relax, show differences in neural responses across different stimulus configurations but sharing the same type of attentional focus. Submatrices off the diagonal (Space-Talker, Space-Relax and Talker-Relax), on the other hand, quantify the dissimilarity in neural responses between different forms of attentional function. (b) Three conceptual model RDMs used to study patterns in EEG RDMs. In Space vs Relax, Talker vs Relax and Space vs Talker, submatrices representing comparisons between attention types are given values of +1s, and submatrices representing comparisons within each attention type are given values of *−*1s. These models can be used to study the representation of attention types.

To study the dynamics of neural representation of attention, we created several conceptual models that capture specific patterns of similarity and differences among conditions. Each ideal RDM model consisted entirely of *±*1s: positive entries for conditions that are very different in the dimension of interest and negative entries for conditions that are very similar (Figure 3b); all other values in the model RDM are irrelevant to comparing particular pair of attentional states and are omitted from further analysis. The “Space vs. Relax” model, for instance, has low dissimilarity within Space and Relax conditions (the Space-Space submatrix in the top left of the RDM and the Relax-Relax submatrix in the bottom right portions of the RDM), and high dissimilarity between these conditions (the Space-Relax submatrix in the bottom left and top right portions of the RDM). All sub-matrices of the model that include a Talker condition are set to zero, effectively excluding them from the analysis, since these conditions give no information about the difference in brain activity patterns for Space and Relax conditions. Similar conceptual models were constructed for “Talker vs. Relax” and “Space vs. Talker” (Figure 3b).

We then compared each subject’s EEG RDMs over time to the ideal model RDMs by computing the correlation between the computed RDM at each time point and the ideal model. Entries with zero values in the ideal model RDMs were first excluded, and the non-zeros entries for both EEG and model RDMs were demeaned before calculating their correlation. For each EEG feature and experimental interval (cue and stimulus period), we tested the correlation between the conceptual model RDMs and the EEG RDMs derived from our data. A high correlation at a time point indicates that the EEG RDM at that time is similar to the model being tested, which implies that there are reliable differences in the brain states during the different attentional states being contrasted. The time series of correlation coefficients from each subject were used as the input for a cluster-based permutation test (CBPT) for statistical inferences.

### 2.7. Statistical analysis

The cluster-based permutation test (CBPT; for details, see (Maris and Oostenveld, 2007)) is a non-parametric statistical test that identifies significant effects that are continuous in the feature space (e.g., in time, space, frequency, etc.). In this study of correlation with conceptual models, we set up the CBPT to identify continuous time intervals in which the average correlation between a given ideal model and the individual RDMs was significantly above zero across subjects. Specifically, we first calculated one-sample t-statistics at each time point to find times in which the mean of the across-subject distribution of the correlation coefficient between an ideal model and the RDM was greater than zero. We then looked for time-clusters across which such significant differences were consistent and persisted longer than would be expected by chance, using a permutation method. We set the cluster formation alpha to 0.05, which led to a series of above-threshold t-values. These t-values formed clusters that were continuous in time, and we used the sum of t-values within each cluster to generate cluster-level test statistics. We then derived a null distribution through randomly permuting data labels. For a one-sample t-test, this permutation process could be achieved by negating the sign of data for a randomly selected subset of all subjects. Using the permutation dataset, we followed identical procedures to generate null-model cluster-level test statistics. Specifically, we again conducted a one-sample t-test at each time point, and calculated the sum of t-values for each found cluster. Only the maximum cluster-level test statistics from each permutation was used to form the null distribution. We repeated this permutation process 10,000 times, which yielded a null distribution with 10,000 cluster-level test statistics. Then, we compared the observed cluster-level test statistics for the actual data against the null distribution, and assigned a p-value to each cluster — this p-value equaled the proportion of the null distribution that were greater than the observed test statistics of the cluster. Any cluster with a p-value less than 0.05 was deemed as significantly above zero in this study.

## 3. Results

Our experiment had multiple conditions that required listeners to adopt different attention strategies and thus to enter different cognitive states. We recorded EEG from 30 subjects while they performed different forms of an attention task using similar stimuli, then extracted information about neural representations from the EEG Scalp Voltage or its Alpha Power. We explored these representations using the difference in neural activity between each pair of conditions, estimated by a linear SVM’s ability to separate them. The average classification accuracy of the SVM provides a measure of the dissimilarity between two conditions at a given time point. We created time-varying representational dissimilarity matrices (RDMs) from all pair-wise dissimilarities across conditions (combinations of stimuli and attentional instructions). Each point (i, j) of an RDM at a given time gives the dissimilarity (i.e., the classifier accuracy) between conditions i and j. We generated two sets of RDMs as a function of time, one with classification values from the EEG Scalp Voltage measured across 64 channels and one with classification using Alpha Power distribution across the scalp.

### 3.1. Representational dissimilarity matrices

#### 3.1.1. Decoding with EEG Scalp Voltage

Figure 4a (top row) shows some of the EEG RDMs derived from the EEG Scalp Voltage during the cue period. Each row or column in an RDM corresponds to a condition index, and these conditions are ordered as shown in Table 1: conditions 1 – 8 are for spatial attention, 9 – 14 are for talker attention, and 15 – 21 are for passive listening. To better visualize information described by the RDMs, we applied multi-dimensional scaling to the dissimilarities entered in the RDM submatrices. In the resulting MDS plots, Euclidean distance between data points represents the dissimilarity between the corresponding conditions (see bottom rows of Figure 4a and Figure 4b).

**Figure 4:**
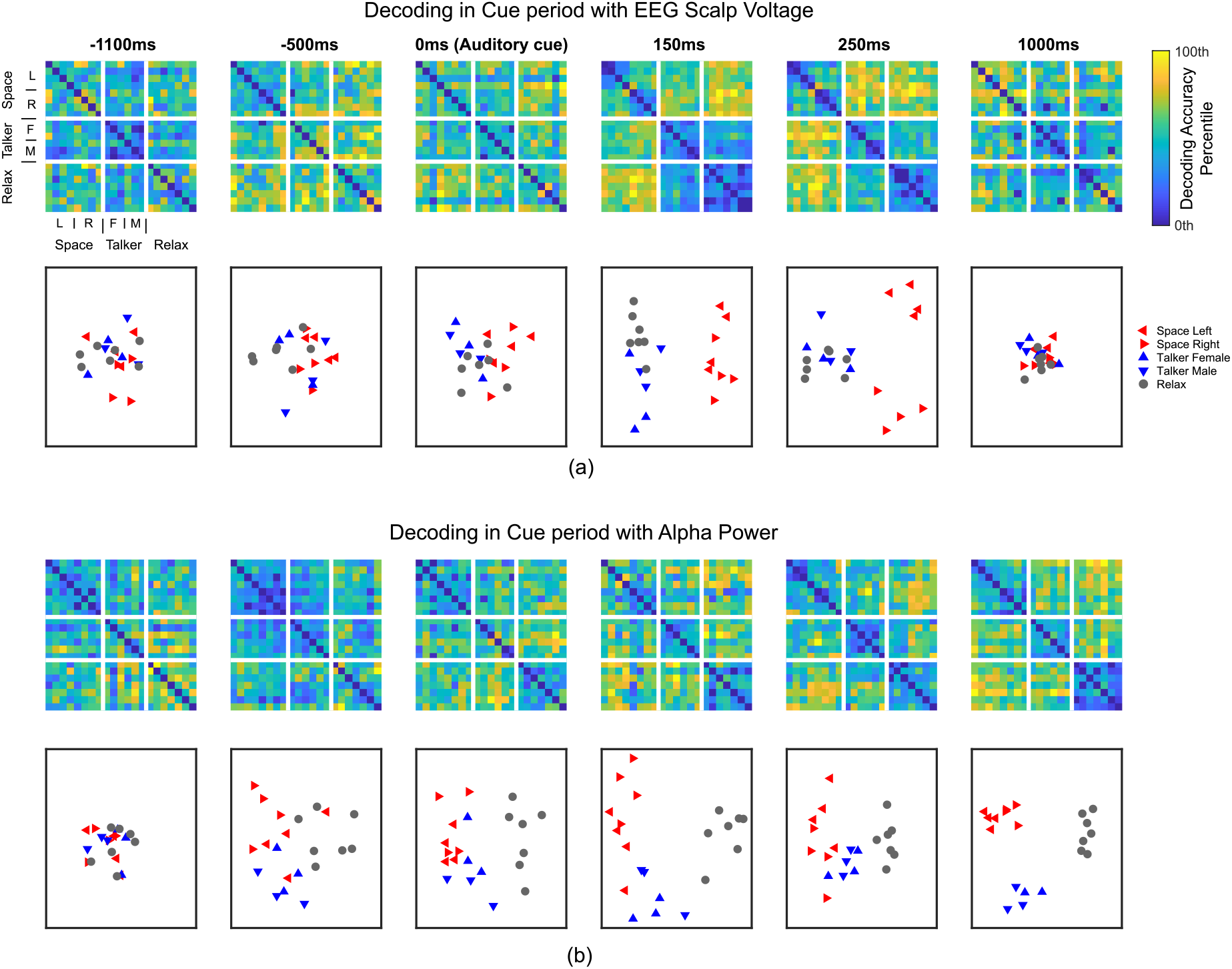
Examples of representational dissimilarity matrix in the cue period, derived from (a) EEG Scalp Voltage, and (b) Alpha Power. −1000 ms denotes the onset of visual cue. 0 ms denotes the onset of auditory cue. Color map was scaled to the 0th and 100th percentile value within each RDM.

To better visualize the patterns in the RDMs, we applied multi-dimensional scaling, averaging dissimilarities across cells within submatrices corresponding to comparisons across attentional condition (e.g., Space-Space, Space-Talker, Space-Relax, etc.). The resulting plots show the average dissimilarity across the sets of conditions. Before the auditory cue (AC) was given, the RDMs do not show a specific pattern. When visualized using MDS, conditions are randomly positioned and intermingled at the start of the cue period (Cichy et al., 2014). Approximately 150 ms after presentation of the AC that gives the specific target for each trial (the target direction for Spatial attention or the target voice for talker attention; see Figure 1), the RDM shows a clear pattern: spatial attention could be decoded accurately from talker and passive listening in feature space, as indicated by the high decoding accuracies in the Space-Relax and Space-Talker submatrices. This separation is clearly visible in the corresponding MDS plot, where the red triangles, representing Space conditions, form a unique cluster. At 250 ms after the AC onset, left attention conditions can be further decoded from right attention conditions. However, these clear representations of attention are transient in EEG Scalp Voltage. The RDMs return to their random pattern towards the end of the cue period (e.g. at 1000 ms).

#### 3.1.2. Decoding with Alpha Power

Information in the alpha band persists beyond that in the EEG Scalp Voltage. At −1100 ms and −500 ms, the RDMs do not form a clear pattern (i.e., after the VC but before the AC), and the corresponding MDS plots show a random scattering of conditions (Figure 4b). At 0 ms and beyond, the Relax conditions could be decoded from the active attention conditions. This is evidenced by the relatively higher decoding accuracies in the Space-Relax and Talker-Relax submatrices than those in the Relax-Relax submatrix in these RDMs. At 1000 ms after the AC onset, Space, Talker, and Relax conditions each form a unique cluster in the MDS plot, indicating that the information in the scalp distribution of alpha oscillation is distinguishable among the three types of auditory attention during this preparatory period.

### 3.2. Correlation with conceptual model RDMs

#### 3.2.1. EEG Scalp Voltage RDMs

We analyzed the similarity between EEG RDMs and three conceptual model RDMs that captured key differences among conditions (Space vs Relax, Talker vs Relax, and Space vs Talker; see Figure 3b). Each model RDM represents the pattern expected if two different types of attention are perfected distinguishable; the correlation between each ideal model RMD with the EEG RDMs can reveal the temporal dynamics of attentional state during a trial.

Figure 5a plots the correlations between conceptual models and the series of EEG Scalp Voltage RDMs. The correlations are significantly above zero only during time windows shortly after a cue, either visual or auditory, is presented. The correlation values following the visual cue are quite weak (correlations of roughly 0.1 or lower), while those following the auditory cue are stronger (peaks greater than 0.2). Figure 5b shows the grand average event-related potentials (ERPs) from channel FCz for each attention type. We selected FCz because the signal in this channel shows the highest contrast among different attention types. By inspecting the ERP, we hope to understand what neural activity features contribute to the neural decoding, and may thus explain some differences between RSA and conventional EEG analyses.

**Figure 5:**
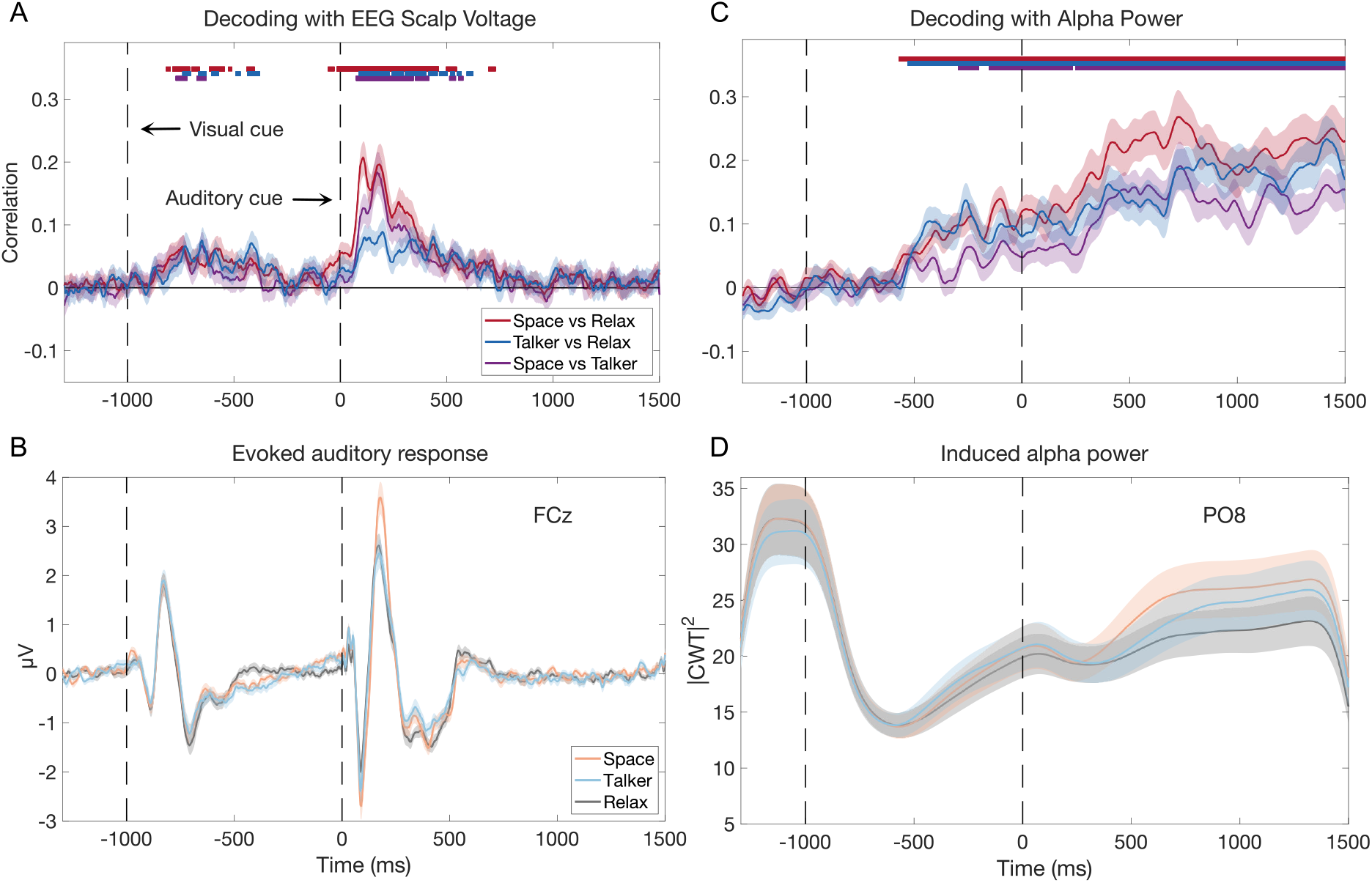
(A) Correlation between the EEG Scalp Voltage RDMs and three conceptual RDMs. (B) The evoked auditory event-related potential at the channel FCz. (C) Correlation between the Alpha Power RDMs and three conceptual RDMs. (D) The induced alpha power at the channel PO8. In all panels, the shaded areas denote the standard error of the mean, across subjects. The bars above the correlation traces denote the temporal clusters in which the model correlation is significantly above zero (determined via a cluster-based permutation test, see Methods 2.7, which the three colors denoting the different condition pairs.

Following the onset of the VC (−1000 ms), we observe a clear ERP waveform. Though the correlations with conceptual models are significantly above zero between approximately −800 ms and −500 ms, the FCz ERPs during this period do not differ greatly between attention types. Following the AC onset (0 ms), however, we observe both significant correlations with the conceptual models and a divergence in ERPs across attention types; the N1 and P2 components at FCz are amplified in Space compared to Talker and Relax conditions, which may have been the driving force behind the Scalp Voltage RDM’s strong correlation with Space vs Relax and Space vs Talker model RDMs between approximately 100 ms and 250 ms. The P1 component is also enhanced in Talker compared to Relax conditions. After the offset of the AC (500 ms), both ERP and the correlation traces return to their baseline.

#### 3.2.2. Alpha Power RDMs

The correlations between Alpha Power RDMs and conceptual model RDMs show a very different pattern. The correlations rise above baseline at around −500 ms (between VC and AC onsets), but then remain significantly above zero throughout the entire cue period (Figure 5c). The correlations for all three conceptual models have very similar trends, although the correlation with the Space vs Talker model takes longer than the other two models to rise to a level that significantly exceeds zero. Figure 5d shows the induced alpha power change at a representative channel, PO8, during the cue period. At this single channel, we do not observe substantial difference between attention conditions until approximately 500 ms after the AC onset, when the Space and Talker show stronger alpha power than the Relax condition.

## 4. Discussion

In this work, we used RSA to demonstrate the feasibility of decoding the representation of auditory selective attention state in EEG signals. We designed a condition-rich experiment that used similar stimuli across conditions but that required listeners to adopt different listening strategies and cognitive states in different task conditions. We recorded EEG from 30 participants while they performed an auditory attention task and, via a neural decoding approach, studied the representation of attention in both the evoked EEG Scalp Voltage and Alpha Power (generalized across different stimuli). We estimated RDMs at each time point and assessed their correlation with conceptual model RDMs that reflect perfect distinctions between pairs of attentional states. The Scalp Voltage of these correlations reveals the brain’s information dynamics during auditory attention.

### 4.1. Application of RSA in studying internal states

The RSA approach to study neural representation (Kriegeskorte et al., 2008) has been applied in multiple domains in neuroscience research, including visual object recognition (Connolly et al., 2012; Cichy et al., 2014, 2016), visual object processing (Kaneshiro et al., 2015), scene perception (Cichy and Teng, 2017), audiovisual integration (Cecere et al., 2017), and semantic categorization of sound (Lowe et al., 2020). In these studies, researchers presented large sets of different visual or auditory stimuli to participants and studied neural representation of the object categories (e.g., human faces, animate objects, etc.) from their EEG, MEG, or fMRI signals. In other words, these studies applied RSA to decode how the brain processes certain properties of the stimulus or stimulus categories, generalized across different stimulus instances. In contrast, in our study we deployed RSA to investigate the internal states of the brain that reflect different forms of attentional focus and executive control. In our study, the auditory stimuli presented in different conditions were either identical or almost identical; what varied was the attentional state of listeners across different tasks: spatial attention, non-spatial attention, and passive listening. The measure of representation in RSA, RDM, characterizes the information encoded in different brain states.

To our knowledge, only one prior study has adopted a similar strategy to study attentional states. Salmela et al. (2016) conducted an RSA study to investigate the dynamics of the fronto-parietal attention network during an audiovisual attention task. They computed RDMs from the EEG Scalp Voltage, and compared them with RDMs calculated from fMRI signals to track the spatiotemporal dynamics of attentional control. One challenge in that study, as in many other RSA studies, is the trade-off between the number of conditions and the number of trials per condition — the experiment becomes impractical if both are large. As a result, the researchers in that study adopted a relatively small number for both (20 trials per condition for 6 conditions, and 5 trials per condition for the other 12 conditions).

Here, we used short stimuli to help manage the trial length so that we could accommodate both more conditions (n = 21) and more trials per condition (n = 36). This can potentially improve the stability and accuracy of neural decoding (Dash et al., 2019), and allow one to look for brain responses that generalize across a range of different stimuli while sharing similar attention focus. Additionally, with a relatively large number of trials per condition, we could afford to estimate representational dissimilarities with cross-validation, which is highly recommended as noise in the dataset can make EEG signals appear to be more dissimilar than they are in reality (Walther et al., 2016; Popal et al., 2019).

### 4.2. Representation of attention in evoked EEG responses

The correlation between EEG RDMs and the ideal conceptual model RDMs reveals time intervals during which particular attentional states have strong representation in EEG signals. When using EEG Scalp Voltage, i.e., the auditory evoked response, for decoding, we identified two intervals in the cue period that are significantly correlated with the attention-type RDMs (Figure 5a).

The first interval ranges from approximately 250 ms to 700 ms after the onset of visual cue (−750 to −300 ms with respect to the auditory cue). This likely reflects information in the response evoked by the visual cue (the words “Space”, “Talker”, and “Relax”) and may represent how the brain encodes information based on the cue received. Information that correlates with differences in attentional state emerges at times that correspond to the late visual response (200 - 800 ms) rather than the early response (*<*200 ms), suggesting that the information encoded during this interval is not driven by some low-level properties (e.g., intensity or location) of the cue, but rather by changes in brain state once the meaning of this cue has begun to be extracted. This may represent the state of initial preparation for attending to the upcoming auditory cues during the cue period.

Following this initial, weak representation of attention (judging from the amplitude of correlation), two more prominent correlation peaks appear in Space vs Relax and Space vs Talker models at approximately 100 ms and 180 ms after the onset of auditory cue, respectively (Figure 5a). The RDMs during this interval show that spatial attention conditions can be differentiated robustly from the other two attention types (Figure 4a); the corresponding multi-dimensional scaling (MDS) plots in Figure 4a also show a clear separation of spatial attention conditions (the red triangles) from the other conditions. Unlike the first interval, the high correlation between neural and model RDMs in this later interval is driven by early auditory evoked responses; the peaks are aligned with the timing of N1 and P2 components in a canonical auditory ERP. This might be due to the fact that the auditory cue (AC) in Space conditions comes from either the left or the right, while the AC in the other conditions comes from the center; in other words, differences in the physical properties of this cue may drive the differences in responses between these conditions. Additionally, we also observe a relatively small, but significant peak in Talker vs Relax during the same time window (Figure 5a). This peak is likely the product of attention modulation rather than the property of the sound, as the ACs in Talker and Relax conditions share the same location. The significant correlation could be caused by the gating effect of attention on sensory inputs, where an attended stimulus evokes greater brain responses compared to an ignored stimulus. Such attention modulation of evoked response has been reported in multiple auditory (Choi et al., 2013; Giuliano et al., 2014) and visual (Bonacci et al., 2020; Hong et al., 2020) attention studies.

### 4.3. Representation of attention in Alpha Power

The correlation between conceptual model RDMs and Alpha Power RDMs in the cue period exhibit a persistent temporal pattern — the correlation with Space vs Relax and Talker vs Relax reaches statistical significance at approximately −500 ms (500 ms after the onset of VC) and stays significantly above zero until the end of the cue period (Figure 5c). Significant correlation between the Alpha Power RDMs and the Space vs Talker model RDM, however, happens relatively late — at approximately −250 ms (750 ms after the onset of VC). The MDS plot at 0 ms shows a clear separation between active attention conditions, whether spatial or non-spatial, and passive listening conditions (Figure 4b). These post-VC alpha-band oscillations may thus indicate a neural process of preparation for the auditory cue (AC) that is shared by both spatial and non-spatial attention.

After the AC plays, the three attention types gradually form unique, separable clusters in the MDS plot (Figure 4b). This visualization shows that towards the end of the cue period, the induced brain response encodes the different types of attention. In other words, the brain is in different internal states depending on whether it is preparing for a spatial, non-spatial, or passive listening trial. Compared to the information encoding in the EEG Scalp Voltage, this representation of auditory attention persists longer.

It should be noted that the temporal resolution in the Alpha Power RDMs is lower than that in the EEG Scalp Voltage RDMs. This is due to the fact that the continuous wavelet transform (CWT) filters input signals with a time window whose length depends on the frequency of interest. In our analysis, the lowest alpha component, evaluated at 8.1 Hz, has a half width at half maximum of approximately 150 ms. Information from future time points, therefore, impacts the RDM calculation at the present time point, and we should be cautious when interpreting the exact time at which information can be decoded from the Alpha Power distribution on the scalp. For example, significant correlation in the Space vs Talker model happens approximately 250 ms before the onset of AC (Figure 4b). We speculate that changes in alpha power distribution *after* the AC onset may contribute to these results. Since each strategy to correct for this causality issue (e.g., downsampling or time shifting) introduces other problems, we choose to present our results as they are; however, we cannot stress enough that the times at which information is present in the distribution of Alpha Power cannot be determined precisely.

### 4.4. Compare RSA to classical EEG analysis

RSA is a powerful tool to study neural correlates within a condition-rich experimental design. The approach allows one to look for neural representations of information that generalize across different conditions (such as trials with different stimuli, as in this study) but that are consistent in some other way (such as the form of attentional focus, as in this study, or the category of an input, as in many past studies).

Classical EEG studies, such as ERP analysis or time-frequency analysis with a binary contrast, typically directly compare the signal or some measures calculated from the signal (e.g., alpha power) between two conditions or condition groups. Instead, RSA concerns the representation — the dissimilarity structure among all conditions — and operates on the *unsigned* difference rather than the signed difference between conditions. This gives it more sensitivity to detect information in certain scenarios, such as when the polarity of EEG differences is inconsistent across participants. For example, one listener may perceive a female voice as more salient than a male voice, and hence show stronger ERPs for a female voice; another listener may have the opposite perception and stronger ERPs for a male voice. In a classical ERP analysis, results from these two listeners would cancel each other out, producing no differences between responses to male and female talkers. In the RSA approach, however, the fact that both the ERPs of both listeners are sensitive to differences in the pitch of talkers means that this information will be retained across these two listeners, because the information encoded in EEG signals is the same, even though the exact representation differs.

The nature of multivariate analysis in RSA also gives it the ability to combine evidence across multiple features, which can be more powerful than a univariate analysis. In Figure 5b and 5d, we show the measures from two electrodes that were used to conduct neural decoding and to construct RDMs. We picked these two electrodes because we observe the greatest differences in their respective measures (i.e., Scalp Voltage or Alpha Power) across types of attention. However, even though the RDMs have significant correlations with conceptual model RDMs at some time windows (Figure 5a and 5c), which hints at there being significant differences across attention types, it does not necessarily translate to significant effects at any single electrode. This is especially true in the induced alpha power at PO8 (Figure 5d) — the correlations become significant before 0 ms, indicating strong information encoding even before the AC was played, while the alpha power at PO8 at 0 ms is hardly distinguishable between attention types. Therefore, compared to some classical approaches, RSA has more sensitivity and less bias as no electrodes need to be pre-selected to define regions of interest, which is sometimes subjective and inaccurate.

### 4.5. Limitations

In this study, we tested the representation of a set of conceptual models in EEG through their correlation with EEG RDMs. These tests were conducted separately for each individual model, so that we could examine one aspect of attentional control at a time. For example, correlating EEG RDMs with the Space vs Relax model RDM only revealed time points in which EEG signals show differences between spatial attention and passive listening. As simple and straightforward as it is, this approach does not account for the fact that our brain is a multi-tasking machine — the cognitive functions encoded by these conceptual models may in fact happen simultaneously. One way to acknowledge the interaction between these functions during the task and examine them in a more integrative manner is to conduct a multi-model fitting analysis (Freund et al., 2021). We can treat the conceptual models in Figure 3b as competing models during the task, and fit the observed EEG RDMs as a linear combination of the conceptual models at each time point. This process will yield a set of model coefficients (or weights) as a function of time, which can implicate the dynamics of cognitive functions encoded by these models, and offer a rigorous framework to compare their relative strength at a certain time point. Another limitation of this study is the relatively small number of conditions in our experimental design compared to previous RSA studies (Cichy et al., 2014, 2016). Using a smaller number of conditions can increase the risk of false discoveries. We mitigated this risk by adopting a non-parametric statistical method (the cluster-based permutation test), which makes no assumptions about noise characteristics. Instead, null distributions are generated from mislabeled data taken from the same experiment – and using the same analysis pipeline – to ensure temporal structure in the data does not bias results. Then statistical inferences are made on a cluster level. We also recruited a relatively large number of participants (n=30) to increase statistical power and lead to smaller bias in our estimation of population means. However, a greater number of conditions, trials per condition, and sample size would have achieved even better power.

Finally, our design was constrained by the use of an attention task, requiring subjects to respond in order to ensure they stayed on task. Most previous RSA studies focused on decoding stimulus categories or traits, rather than internal processing states, which freed them from using active tasks requiring subject responses. Future studies should consider designing experiments in such a way that multiple trials and/or conditions could be acquired per subject response. Such an approach would reduce the time needed to acquire a dataset that is ideal in size (i.e., number of conditions and trials per condition) for an RSA study.

## 5. Conclusions

This paper demonstrates that the representational similarity analysis framework can be used to investigate the neural representation of executive function (specifically attentional state). We designed a condition-rich experiment and recorded EEG signals while listeners participated in an auditory attention task. We extracted representational dissimilarity features from the EEG Scalp Voltage and Alpha Power. Comparison of the resulting neural representation (summarized by dissimilarity matrices that contrast all possible conditions within a rich set of trial types) with ideal conceptual models confirms the feasibility of using RSA to investigate internal cognitive state. Specifically, we identified time intervals in which particular contrasts in attentional state, such as the difference between attention types or between attention to different locations, emerge in the neural responses measured from the EEG Scalp Voltage and the Alpha Power distribution on the scalp.

## Declaration of competing interest

The authors declare no financial conflicts of interest.

## Acknowledgements

This study was supported by the the Office of Naval Research [Project number N00014-18-1-2069].

## Notes

### Competing Interest Statement

The authors have declared no competing interest.

